# Warmer temperatures favor slower-growing bacteria in natural marine communities

**DOI:** 10.1101/2022.07.13.499956

**Authors:** Clare I. Abreu, Martina Dal Bello, Carina Bunse, Jarone Pinhassi, Jeff Gore

**Affiliations:** Physics of Living Systems, Department of Physics, Massachusetts Institute of Technology; Cambridge MA, USA; Department of Biology, Stanford University; Stanford CA, USA; Helmholtz Institute for Functional Marine Biodiversity at the University of Oldenburg; Oldenburg, Germany; Centre for Ecology and Evolution of Microbial Model Systems, Department of Biology and Environmental Science, Linnaeus University; Kalmar, Sweden

**Author notes:** These authors contributed equally to this work (Clare I. Abreu, Martina Dal Bello). Department of Marine Sciences, University of Gothenburg; Gothenburg, Sweden (Carina Bunse).

## Abstract

Earth’s life-sustaining oceans harbor diverse bacterial communities that display varying composition across time and space. While particular patterns of variation have been linked to a range of factors, unifying rules are lacking, preventing the prediction of future changes. Here, analyzing the distribution of fast- and slow-growing bacteria in ocean datasets spanning seasons, latitude, and depth, we show that higher seawater temperatures universally favor slower-growing taxa, in agreement with theoretical predictions. Our results explain why slow growers dominate at the ocean surface, during summer, and near the tropics, and provide a framework to understand how bacterial communities will change in a warmer world.

## Main Text

Oceans cover 70% of the surface of our planet and are home to myriad bacterial species that are responsible for many of Earth’s crucial biogeochemical functions (*1, 2*), including fixing carbon and nitrogen, recycling nutrients and dissolved organic matter, and degrading biomass. Surveys of the ocean microbiome over broad spatial and temporal scales have revealed repeatable patterns across seasons (*3–6*), latitude (*7*), and depth (*8*), which have been linked to a variety of environmental factors, including temperature (*3, 4, 8, 9*), water mixing (*10, 11*), nutrient availability (*12*), light (*13*), and the occurrence of phytoplankton blooms (*6*). Nevertheless, general principles underlying the compositional turnover of vital marine bacterial communities are lacking, impairing our ability to predict the response of ocean systems to future environmental changes.

The distribution of fast- and slow-growing taxa is a powerful trait-based description of the structure of a bacterial community (*14, 15*), just as it is for multicellular organisms ranging from plants (*16*) to corals (*17, 18*). Maximum growth rates are a key component of the diverse life history strategies exhibited by micro- and macro-organisms (*19*–*23*), and determine the ability of a species to survive and compete in a given environment (*24*). In aquatic bacteria, maximum growth rate differentiates fast-growing copiotrophs, found in nutrient-rich waters, from slow-growing but efficient oligotrophs, which can grow in nutrient-poor environments (*25*–*29*). While the role of nutrients in the distribution of fast- and slow-growing bacteria has received a lot of attention (*12, 30*–*32*), the effects of temperature have been comparatively less explored, despite ample evidence that temperature is a fundamental driver of marine communities (*3, 4, 8, 33*).

To explore the distribution of fast- and slow-growing bacteria along principal axes of temperature variation in the ocean, we gathered seven 16S rRNA gene amplicon sequencing datasets of marine bacterial communities encompassing wide seasonal, latitudinal, and depth gradients. Given that the maximum growth rate of a bacterial species is approximately proportional to its ribosomal RNA operon copy number (*34, 35*), we inferred maximum growth rates of bacterial community members in these datasets by matching observed taxa to the Ribosomal RNA Database (rrnDB *36*). We summarized the distribution of fast and slow growers in a community with its abundance-weighted mean copy number (MCN), which represents the expected rRNA copy number of a randomly sampled cell (see Materials and Methods). A large MCN indicates greater relative abundances of fast-growing bacteria, while more abundant slow-growing bacteria drive the MCN to smaller values. In Fig. 1, we focus on three datasets of free-living bacteria (the fraction of the biomass that passes through a 3*μ*m filter and is deposited onto a 0.2*μ*m filter) in which temperature changes either seasonally (Linnaeus Microbial Observatory (LMO) in the Baltic Sea, green dot in Fig. 1A), across ~100 degrees of latitude (ANT 28-5 cruise (*37*–*39*), yellow dots), or across a depth of 1,000 meters (TARA Oceans project (*8*), purple dots).

**Fig. 1.**
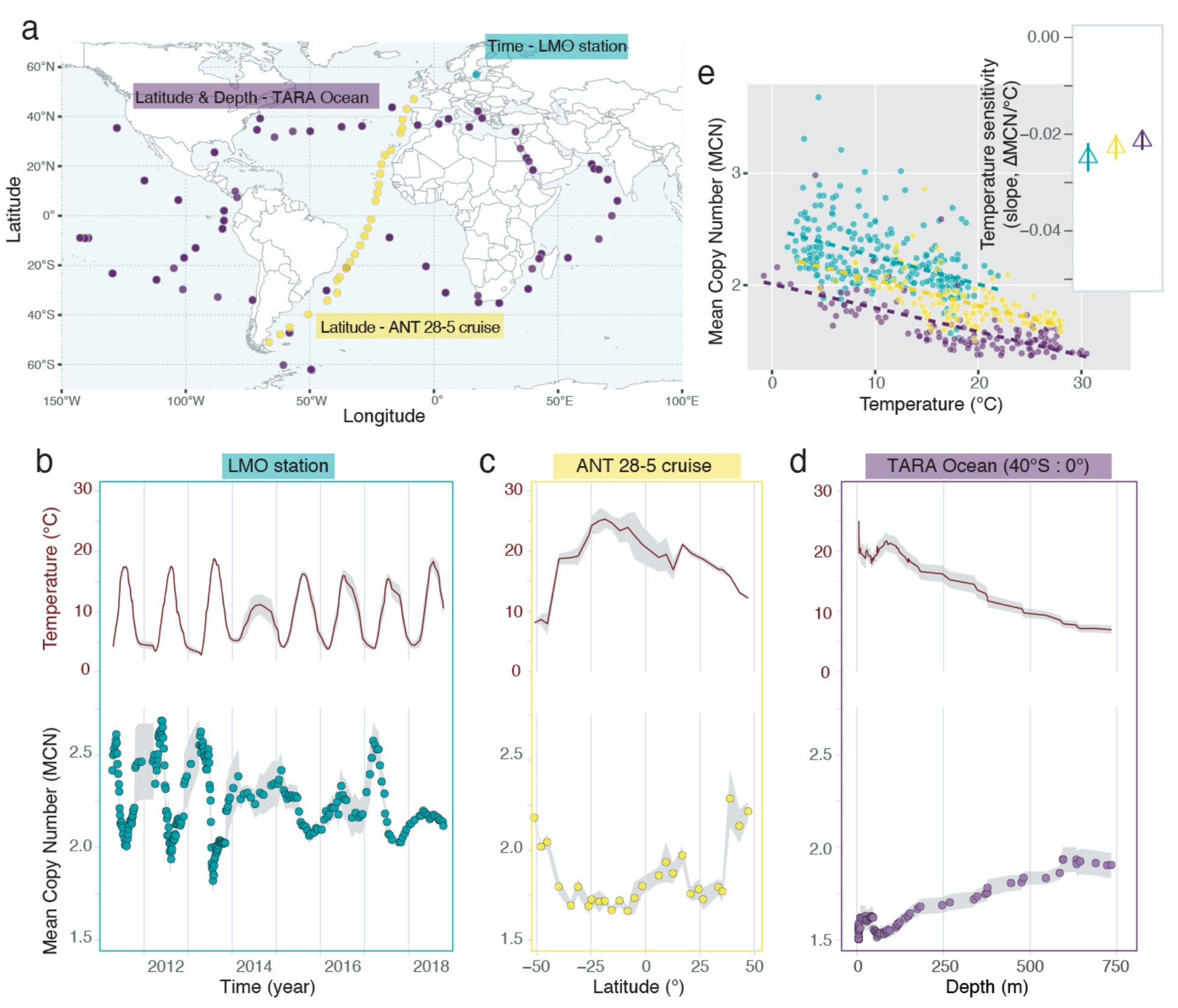
Slower-growing taxa dominate during summer, at low latitudes and at the surface of the ocean. (**A**) Sampling locations of the Linnaeus Microbial Observatory (LMO, green), the ANT 28-5 cruise (yellow), and the TARA Ocean Project (purple) span several seasons, latitudes and depths. (**B**) Temporal trajectories of temperature and weighted mean copy number (MCN) values for the LMO station (eight years, with denser sampling in the first three years and monthly sampling starting in 2014). Each point in the MCN panel is the rolling average ± SE (shaded grey ribbon) with span = 7 observations. Same for temperature but only the trajectory is shown. (**C**) Latitudinal variation in temperature and MCN in the ANT cruise. Each point in the MCN panel is the average across 5 sampling depths (0-25, 25-50, 50-80, 85-120, 120-200m) ± SE (shaded grey ribbon). Same for temperature but only the trajectory is shown. (**D**) Variation in depth of temperature and MCN in the samples collected between 0 and 40°S of the TARA Ocean project. Each point in the MCN panel is the rolling average ± SE (shaded grey ribbon) with span = 19 observations. Same for temperature but only the trajectory is shown. (**E**) MCN is significantly negatively correlated with temperature across the three datasets (temperature sensitivities in the inset: green, LMO = −0.025 ± 0.003 ΔMCN/°C (*p* = 3.78e-15), yellow, ANT = −0.023 ± 0.002 ΔMCN/°C (*p* = 6.83e-15), purple, TARA = −0.021 ± 0.002 ΔMCN/°C (*p* < 2e-16)). The colored dashed lines are linear regressors.

We found striking and consistent changes in the distribution of fast- and slow-growing taxa associated with temperature variation over seasons, latitude, and depth. The MCN of LMO free-living bacterial communities showed recurring seasonal fluctuations, with the lowest values observed during summer when temperatures were highest (Fig. 1B). The MCN of free-living communities along the Atlantic transect was higher at higher latitudes and decreased towards the equator, mirroring the latitudinal variations in temperature (Fig. 1C). In the TARA Ocean data, we focused on samples between the equator and 40°S to minimize latitudinal variation and assess how the distribution of growth rates would change with depth. We found that MCN increased as depth increased and temperature decreased (Fig. 1D). In all three cases, higher temperatures were associated with an increase in the relative abundance of slow-growing taxa with low copy number.

Temperature sensitivity, i.e.: the slope of the regression between MCN and temperature, is consistent across the three datasets, revealing a significant decrease of the mean copy number within the community by ~0.5 over the range of temperature observed in each dataset (LMO temperature sensitivity = −0.025 ± 0.003 ΔMCN/°C (*p* = 3.78e-15), ANT = −0.023 ± 0.002 ΔMCN/°C (*p* = 6.83e-15), TARA = −0.021 ± 0.002 ΔMCN/°C (*p* < 2e-16); Fig. 1E). For example, the MCN of the LMO dataset spans between ~2.5 at the lowest temperature and ~2 at the highest (Fig. 1E). Our study suggests that slow growers are relatively more abundant where and when temperatures are higher, i.e.: at the surface of the ocean, during boreal summer, and around equatorial and tropical regions, and that this direct effect of temperature is a general feature of the ocean microbiome.

We previously demonstrated that in a two-species laboratory community of soil-derived bacteria, the slower-growing species usually increases in relative abundance at higher temperatures (*40*). Here, we extend the two-species Lotka-Volterra model presented there (*40* but see also fig. S1-S3 and Supplementary Text for derivation) to a simulated community of 100 species (Fig. 2A). In this generalized Lotka-Volterra (GLV) model, all species’ growth rates are proportional to their rRNA copy number and increase with temperature (Fig. 2a, fig. S1; see Materials and Methods and Supplementary Text). When subjected to a global mortality rate approximating the effect of grazing, viral lysis, and senescence in the oceans (*41–44*), the mean simulated MCN decreases with temperature (Fig. 2B and fig. S4), equivalent to a mean reduction of ~0.7 per ~30-degree increase in temperature.

**Fig. 2.**
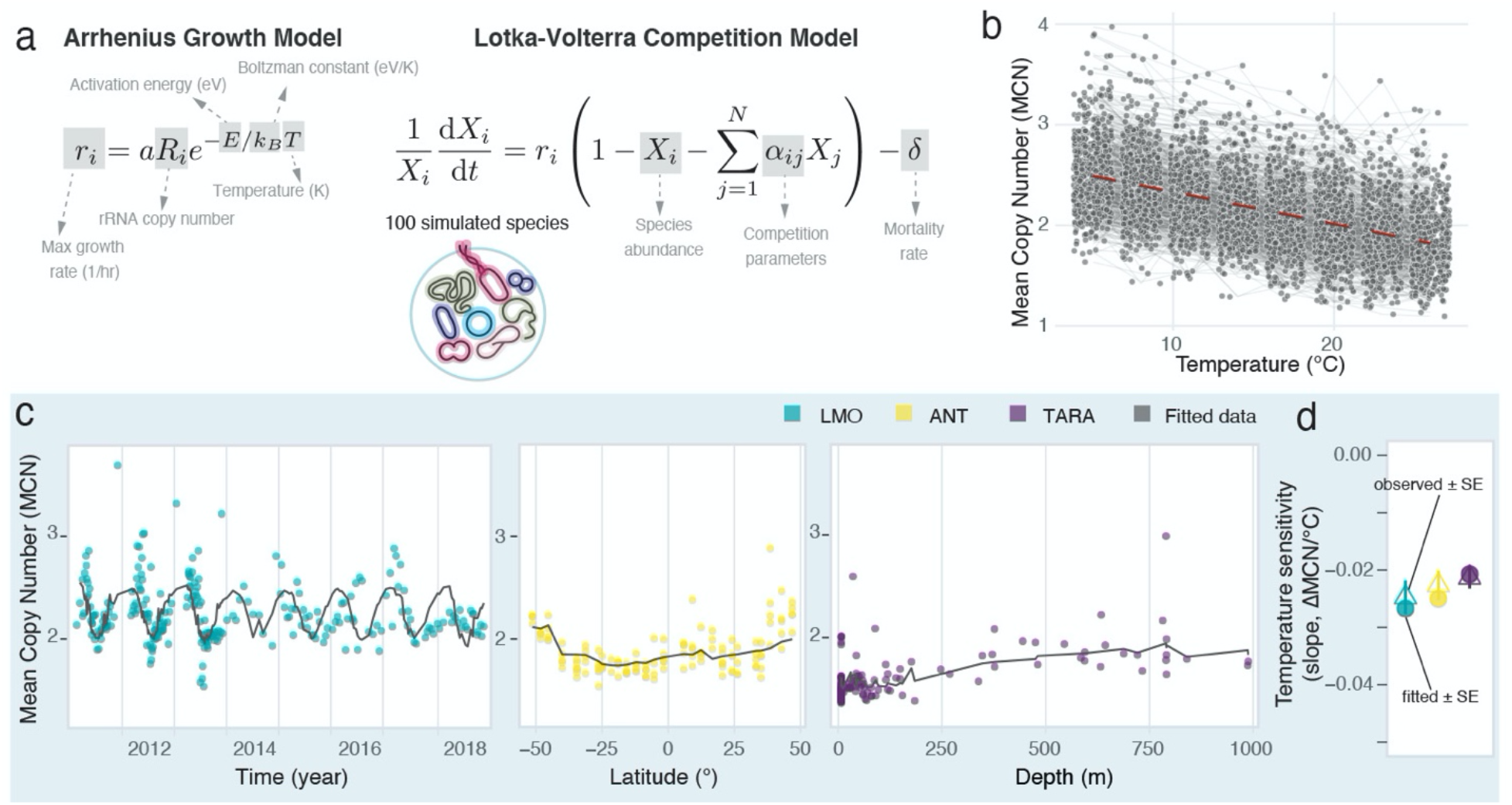
Observed mean copy number is well-fit by an inter-species Lotka-Volterra competition model. (**A**) The growth of a simulated species is proportional to its rRNA copy number and temperature (in K) according to the Arrhenius Model, which we plug into the Lotka-Volterra competition model with a community-wide mortality rate, *δ.* rRNA copy numbers are drawn from a geometric distribution, (1 – *p*)^*k*-1^ *p*, where *k* is the copy number and the parameter *p* accounts for the fraction of simulated species with copy number = 1 in the community starting distribution. (**B**) 500 trajectories of MCN of simulated 100-species communities as a function of temperature. (**C**) Observed (colored dots, green LMO, yellow ANT, purple TARA) and fitted (black lines) weighted mean copy number (MCN) values across years (LMO), latitude (ANT), and depth (TARA, samples between the equator and 40°S). (**D**) Temperature sensitivity (ΔMCN/°C) calculated on fitted data (full circles, slope ± SE) compared to observed temperature sensitivities (open triangles, slope ± SE).

We then used the model to fit observed temperature data and assess the quantitative agreement between observed and simulated MCN for each dataset. We found strong agreement between observed and simulated MCN over the temperature ranges reported in the three datasets (Fig. 2C, D). Differences in the fits to the three datasets reflect different fractions of taxa with a single copy number in the starting distribution of the simulated communities (see Materials and Methods and fig. S5). In the case of the LMO data, the observed MCN appeared to lag behind the model, a result that can be seen in a continuous-time simulation with oscillating temperature (fig. S6). Our theoretical analyses emphasize that the generic effect of increased temperature favoring slower-growing taxa may be a dominant force in structuring bacterial communities in the oceans.

The negative correlation between MCN and temperature in the three datasets might depend on other environmental variables, e.g., nutrients (figs. S7-S9). If the negative correlations were spurious, the inclusion of other environmental variables in a multivariate regression model would result in a change in either the sign or the statistical significance of the temperature coefficient. Results of this analysis (see Materials and Methods) reinforced that the distribution of fast and slow growers of LMO, ANT and TARA free-living communities is strongly affected by temperature, which in all three instances exerts a statistically significant negative effect on MCN (Fig. 3 and table S1, parametric coefficient *γ_temp_* = −0.031, *p* < 0.001 LMO, *γ_temp_* = −0.011, *p* < 0.01 ANT, *γ_temp_* = −0.018,μ < 0.001 TARA). In the LMO dataset, however, increases in inorganic nitrogen favored slow growers (Fig. 3 and table S1, parametric coefficients for ammonium *γ*_*NH*_4__+ = −0.082, *p* < 0.05 and nitrate *γ*_NO_3__- = −0.068, *p* < 0.1) while phosphate and DOC (dissolved organic carbon) did not significantly affect MCN temporal dynamics (Fig. 3 and table S1). In the TARA Oceans dataset, more abundant phosphate and nitrate at high latitudes and deeper in the water column favored fast growers (Fig. 3, table S1, parametric coefficient for phosphate *γ*_PO_4_^3-^_ = 0.094, *p* < 0.01), contributing to the observed latitudinal trend in MCN, while oxygen did not have significant impacts (Fig. 3, table S1). While other factors may be important for community compositional turnover (see also the effect of chlorophyll *a* concentration on heterotrophic communities (fig. S10 and table S1)), overall, these results show that temperature, more consistently than any other environmental variable, explains variations in the spatial and temporal distribution of fast- and slow-growing marine bacteria.

Bacteria identified as oligotrophs (slower-growing, energy-efficient taxa), including the cosmopolitan SAR11 clade (*45*), are extremely abundant across the ocean, particularly in surface waters and warm oligotrophic gyres (*46*). This observation raises the question of whether the temperature trend in our study simply mirrors the biogeography of SAR11 and other oligotrophs (*12*). However, excluding taxa with copy number = 1 from the calculation of MCN in the LMO, TARA and ANT datasets did not affect the negative relationship between temperature and MCN (Fig. 3, table S1 and fig. S11). The robustness of the effect of temperature when oligotrophs with copy number = 1 are excluded is confirmed in two additional time series of free-living bacterial communities: a seven-year survey at the Service d’Observation du Laboratoire Arago (SOLA) sampling station in the North Western Mediterranean Sea (*47*), and a five-year study at the San Pedro Ocean Time-series (SPOT) station, ~10 km off the coast of Los Angeles, California (*48*) (Fig. 3, fig. S11, S12). These results indicate that the trend of high temperatures favoring slow growers is not dependent on the biogeography of oligotrophs. Conversely, SAR11 diversity cannot be captured by the MCN: all SAR11 ecotypes (*45*) are categorized together for having a single rRNA copy number, despite their ecological differences.

**Fig. 3.**
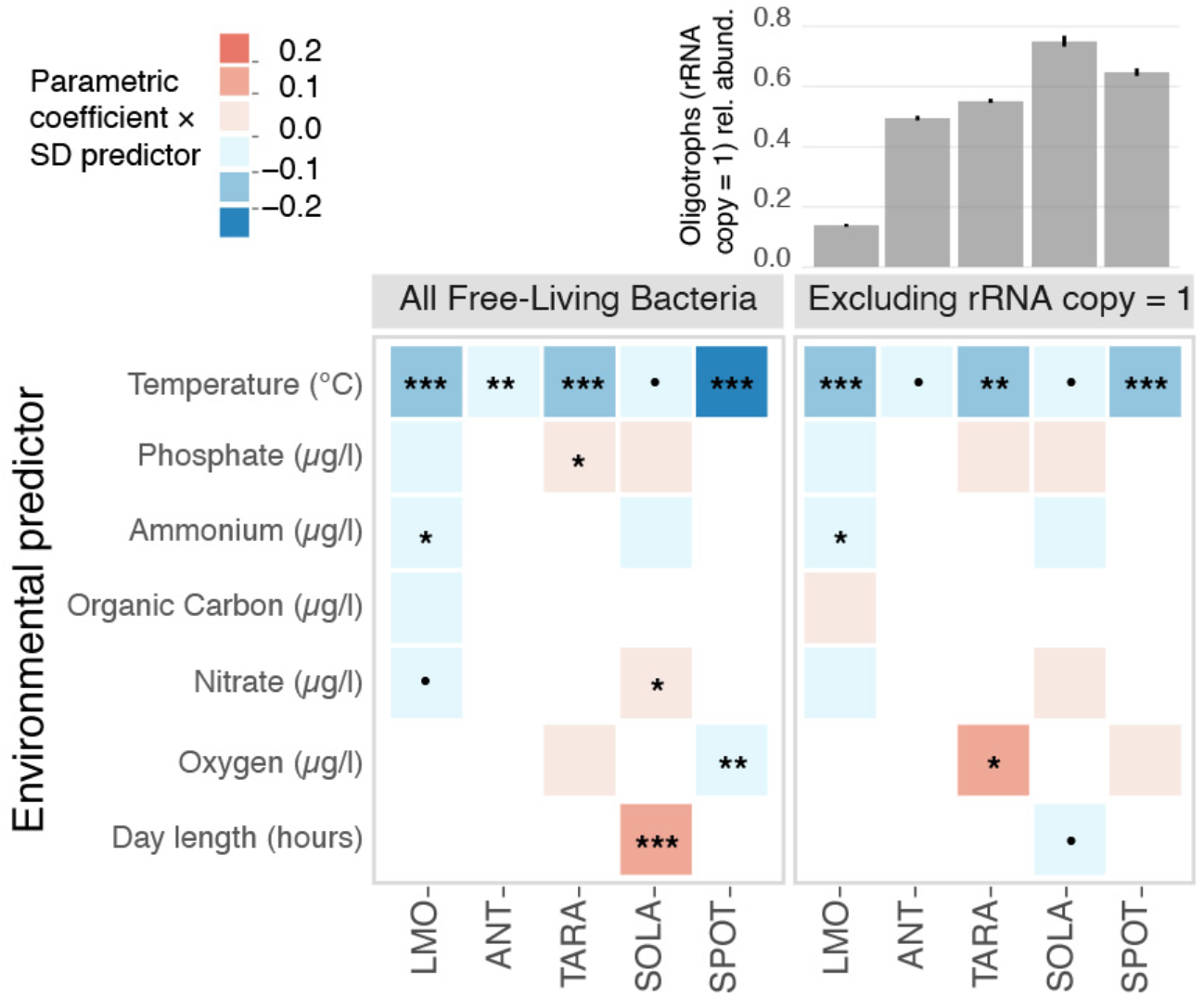
The negative correlation between MCN and temperature is robust to the consideration of other environmental drivers and it is not affected by the removal of oligotrophs. Heatmaps visualize the results of the statical models used to evaluate the effects of environmental predictors on MCN for the three main datasets, LMO, ANT and TARA, and two additional datasets, SOLA and SPOT, all of which include free-living communities (left, all free-living bacteria are considered in the calculation of MCN; right, taxa with rRNA copy number =1 excluded). The color represents the value of the parametric coefficient multiplied by the standard deviation (SD) of the corresponding environmental predictor. Significance codes: ‘***’ p<0.001; ‘**’ p<0.01; ‘*’ p<0.05; ‘.’ P<0.1; ‘ ‘ n.s. The bar plot on the upper right shows the average proportion of oligotrophic bacteria (i.e.: taxa with rRNA copy number =1) in each dataset listed below.

While we have focused on free-living bacteria (as defined by filter fractions between 0.2 and 3 *μ*m), an important question is how temperature affects the dynamics of particle-associated communities (see fig. S13 and table S1), where spatial processes might be important (see *49, 50* for theoretical predictions about the role of nutrient concentrations in spatially extended settings). Another question is whether our conclusions depend on the method for estimating growth rates with rRNA copy number. To test this, we used another method based on codon usage bias (*57*), which yielded an overall negative relationship between temperature and weighted mean growth rate (fig. S14). Altogether, our study does not exclude the role of other processes in the spatial and temporal turnover of the ocean microbiome, but shows that the direct effects of temperature on the distribution of growth strategies have been generally overlooked.

Our proposed rule of higher temperatures favoring slower growers is potentially applicable to any community subject to growth and mortality and could underlie some previously observed patterns. Among common cyanobacteria in the ocean, the slow-growing *Prochlorococcus* dominates near the equator but decreases in abundance towards the poles, where the comparatively fast-growing *Synechococcus* persists (*52*) (see *19, 53, 54* for cyanobacteria growth rate estimates). Similarly, in a 16-year time series of observations of a phytoplankton community at a nearshore site on the Northeast US Shelf, picoeukaryotes with comparatively faster growth rates were outnumbered by *Synechococcus* populations during warmer months(*55*). Finally, in mid-latitude forest habitats, a 20-year-long *in situ* soil warming experiment resulted in a decrease in the MCN of bacterial communities that was attributed to an assumed decline in nutrients (*56, 57*). Interestingly, another study measured a massive reduction in RNA content after incubating a soil bacterial community for one week at increased temperature compared to an ambient control, despite no difference in nutrients between conditions (*58*). The authors attributed the decrease in RNA to an intracellular reduction in ribosome concentration, but a reduction in MCN of the soil community is also consistent with these results. In all of the above examples, we propose that the direct influence of temperature on ecological community composition presents a new interpretation for these observed patterns.

Connecting unifying rules to biogeographical drivers of bacterial community composition throughout the global ocean is an important challenge, especially considering the effects of climate change (*59, 60*). Many studies show that temperature sets the biogeography of marine bacterial species by imposing constraints on their ability to grow, and thereby predict future compositional changes based on the thermal limits of each species (*33*). Our study, comprising a simple model and seven datasets, substantially expands on these predictions by highlighting that, over the broad thermal range of the ocean, temperature has a generic effect on community composition by determining the distribution of fast- and slow-growing taxa. We have presented macroecological patterns in the distribution of growth strategies in bacterial communities across ocean temperature gradients, revealing that slow growers are consistently more abundant around the tropics, during summer, and at the surface of the ocean, in agreement with our theoretical predictions. Overall, our results emphasize that temperature plays a direct role in structuring bacterial community composition and global biogeography of marine bacteria and suggest that warming ocean temperatures may lead to increases in the abundance of slower-growing taxa.

## Materials and Methods

### Datasets

To explore the distribution of bacterial life strategies along principal axes of temperature variation in the ocean, we gathered 16S rRNA gene amplicon sequencing datasets of marine bacterial communities encompassing wide seasonal, latitudinal, and depth gradients.

We focused on three particular datasets in the main text to represent the three principal axes of temperature variation in free-living bacterial communities (Fig. 1a). First, to analyze seasonal data, we used an eight-year pelagic microbial time series from the Linnaeus Microbial Observatory (LMO) in the Baltic Sea, 11 km off the northeast coast of Öland (N 56°55.8540’, E 17°3.6420’) (green dot in Fig. 1a). Seawater samples have been collected since 2011 on a monthly to weekly basis (*5, 6*), together with environmental variables like temperature, inorganic nutrients (nitrate, phosphate and ammonium), dissolved organic carbon (DOC), and chlorophyll *a* concentrations (fig. S7) (*61*). The LMO dataset includes free-living (<3*μ*m and >0.2 *μ*m) as well as particle-attached (>3 *μ*m) and non-fractionated (>0.2 *μ*m) filter fractions. DNA was extracted from filters according to (*62*) and modified after (*63*). We then amplified the V3V4 region of the 16S rRNA gene with the primers 341f-805r (*64*). Amplicon sequencing for LMO data was undertaken at the Science for Life Laboratory, Sweden on the Illumina MiSeq platform (2 × 300 bp paired-end reads). Subsequently, the Ampliseq pipeline (https://github.com/nf-core/ampliseq)(*65*) was applied with DADA2 to infer amplicon sequence variants (ASVs) (*66*). The used bioinformatic software versions were: nf-core/ampliseq = v1.2.0dev; Nextflow = v20.10.0; FastQC = v0.11.8; MultiQC = v1.9; Cutadapt = v2.8; QIIME2 = v2019.10.0. Taxonomic annotation of LMO ASVs derive from the SILVA database (version132) (*67*).

To assess bacterial growth distributions across latitude and depth, we used datasets from two published cruise reports. In 2012, the five-week cruise ANT 28-5 collected seawater samples in the epipelagic zone (20-200m) of 27 stations in the Atlantic Ocean along a transect spanning ~100 degrees of latitude: from the polar regions of South America to the waters off the coast of England (yellow dots in Fig. 1a) (*37–39*). The ANT dataset does not contain publicly available environmental data other than temperature measurements (fig. S8), but does include small (<8*μ*m and >3*μ*m) and large (>8*μ*m) particle-attached in addition to free-living (<3*μ*m and >0.2 *μ*m) filter fractions. The TARA Oceans project was an ambitious four-year expedition (2009-2013) conducted in a modified sailboat, taking samples from 210 globally distributed sites (purple dots in Fig. 1a) at depths from the surface down to 1,000 meters (*68, 69*). All TARA samples are free-living filter fractions (<3*μ*m or <1.6*μ*m and >0.2 *μ*m), and metadata on phosphates, nitrates, and oxygen is included (fig. S9).

In addition to these three datasets, we analyzed three other time series: a three-year study at the Pivers Island Coastal Observatory (PICO) site (34.7181 °N 76.6707 °W) near the Beaufort Inlet (US East Coast) (*4*), a seven-year survey at the Service d’Observation du Laboratoire Arago (SOLA) sampling station (42°31’N, 03°11’E) in the Bay of Banyuls-sur-Mer, North Western Mediterranean Sea, France (*47*), and a five-year study at the USC Microbial Observatory at the San Pedro Ocean Time-series (SPOT) station in the San Pedro Channel (33.55°N, 118.4°W) (*48*). Finally, we analyzed the effects of depth and latitude in the latitudinal P15S GO-SHIP transect, a 7,000-km decadally repeated transect from the ice edge (~66°S) to the equator (0°S) in the South Pacific Ocean (*70*) (fig. S10). Both SOLA and SPOT samples are free-living filter fractions (<3*μ*m and >0.2 *μ*m SOLA, <1*μ*m and >0.2 *μ*m SPOT), while PICO and P15S GO-SHIP samples are non-fractionated (>0.2 *μ*m).

Available environmental variables for PICO include temperature, insolation, nitrate + nitrite, phosphate, ammonium, dissolved inorganic carbon (DIC), and chlorophyll *a* concentrations; for SOLA: temperature, length of the day in hours, nitrate, phosphate, ammonium, and chlorophyll *a* concentrations. Bacterial samples of SPOT were taken at five depths (5, 150, 500, 890 m and at the depth corresponding to the deep chlorophyll maximum, DCM), but only temperature and oxygen measurements were available for all the sampled depths. Particulate organic carbon (POC) and chlorophyll a concentration could be obtained only for the 5m samples, while phosphate and nitrate for 5m and DCM samples. Microbial communities along the P15S GO-SHIP transect have been sampled in 80 stations from the surface to 6000m. Temperature, phosphate, nitrate + nitrite, ammonium, oxygen, and chlorophyll *a* concentration measurements are available from the surface to the mixed layer depth (MLD, around 150m). For samples below the MLD, ammonium and chlorophyll *a* concentration were not available. Following (*70*), we analyzed the data above and below the MLD separately: we used the data within the MLD to explore community growth relationship with environmental variables along the latitudinal gradient and data below the MLD to explore community growth relationship with environmental variables across depths.

### MCN calculation

We used the Ribosomal RNA Operon Copy Number Database (rrnDB *36*) to infer maximum growth rates of bacterial community members in all datasets, because the maximum growth rate of a bacterial species is approximately proportional to its rRNA operon copy number (*34, 35*). The weighted mean copy number (MCN) of a community represents the expected rRNA copy number of a randomly sampled cell in the community and hence is a proxy for the distribution of fast- and slow-growing taxa in a bacterial community. To calculate it, we downloaded the pan-taxa statistics from the *rrnDB* (version 5.6, https://rrndbumms.med.umich.edu/) and matched classified ASVs from each dataset to the listed mean rRNA copy number corresponding to the lowest available rank (for example, if a family-level match was available in the absence of a species- or genus-level match). For each sample in each dataset, we calculated the MCN as an average rRNA copy number of the sample weighted by the relative abundances of each ASV, as explained in Box 1:

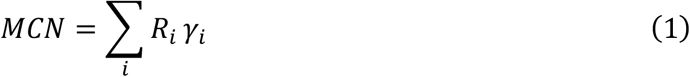

where *γ_i_* and *R_i_* are the relative abundance and assigned copy number of ASV *i*. Since the true relative abundances are obscured by differences in copy number, however, we first divided the sequenced abundances of each ASV, *X_i_*, by their copy numbers, *R_i_*, and normalized to get the true relative abundances:

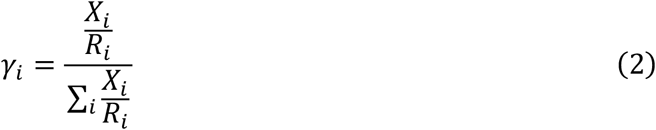

Plugging Equation (1) into Equation (2) gives the MCN in terms of sequenced abundances and assigned copy numbers:

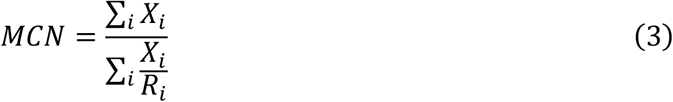

### Models and simulations

In this study, we employed two types of models to describe bacterial community dynamics: generalized Lotka-Volterra equations and a linear consumer-resource model.

In both types of equations, we modeled maximum growth rate as a function of temperature with the Arrhenius model, as shown in Fig. 2A and described in fig. S1:

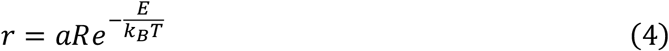

Maximum growth rate *r* of all species increases uniformly with temperature and is proportional to rRNA copy number *R*, which is an integer ranging from 1 to 10. We drew the copy numbers of simulated species from a geometric distribution, (1 – *p*)^*k*-1^ *p*, where *k* is the copy number and the parameter *p* represents the fraction of taxa with copy number = 1 in the starting distribution of the community. We set the spectrum of growth by bounding the maximum possible rate at the highest temperature to known limits (minimal doubling time of ~10 minutes(*51*)), including the prefactor *a* (set to 1.7 * 10^s^ in the fig. S1 simulation). *T* is temperature in Kelvin (range: 278-298), and *E* represents the activation energy (set to 0.33 eV in fig. S1). In the traditional Arrhenius equation, *r* represents the frequency of collisions resulting in a reaction, and has units of 1/time. Here, both growth rate and therefore the prefactor, *a*, have units of 1/hour. Activation energy *E* has units of eV, the same units as *k_B_T* (temperature multiplied by the Boltzmann constant *k_B_*).

The generalized Lotka-Volterra equations are the primary model used in the main text; the details of the linear consumer resource model are available in fig. S3. In the Lotka-Volterra model, all species are subject to an added mortality rate *δ*, across a range of temperatures. The model contains one equation for each species:

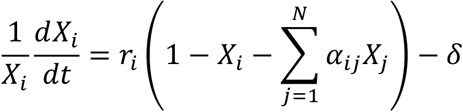

The analytical solution to the two-species model, showing that increasing temperature favors the slower grower, is available in the Supplementary Text. In the two-species example shown in fig. S1, we set *δ*=0.1/hr, *α*_12_ 0.6, *α*_21_=1.3.

In Fig. 2, we simulated 100 interacting species in Python, with competition coefficients *α_ij_* drawn from a normal distribution and rRNA copy numbers drawn from a geometric distribution. In fig. S4, we show that the variance in the MCN-temperature correlation is similar when comparing two cases: 1) if the characteristic parameters of these distributions are randomly drawn for each simulation, and 2) if the characteristic parameters of these distributions are the same for all simulations. Fig. 2B shows the results of case 1), where the mean of the normal distribution of *α_i,j_* was set to 0.5 (<1 allows for coexistence—see Supplementary Text; we also note that the outcome is independent of mean interaction in the two-species solution—see fig. S2) and the standard deviation set to half the mean, or 0.25, the geometric distribution parameter *p* of the rRNA copy numbers was set to 0.8 (in the geometric distribution, (1 – *p*)^*k*–1^ *p, k* is the copy number and *p* represents the fraction of the starting community with copy number = 1), the mortality rate *δ* was set to 0.07, the activation energy *E* was set to 0.33 eV, and the prefactor *a* set to 170,000. In case 2), the mean of the normal distribution of *α_ij_* was randomly drawn from a uniform distribution [0.1, 1] and the standard deviation was set to half the mean, the geometric distribution parameter *p* of the rRNA copy numbers was drawn from a uniform distribution [0.6, 0.9], the mortality rate *δ* was drawn from a uniform distribution [0.03, 0.2](*42*), and the activation energy *E* was drawn from a uniform distribution [0.1 eV, 0.6 eV] (*71, 72*), with the prefactor *a* set to the mean of 0.46e ^*E*/(*k_B_*\*300)^ and 0.05*e*^*E*/(k_B_*,278)^ (ensuring that growth rates with the highest copy number at high temperature did not exceed realistic rates, and that the growth rate of species with copy number = 1 exceeded most mortality rates at the lowest temperatures).

### Fitting model to datasets

To fit the model to the datasets, we simulated the 100-species Lotka-Volterra competition equations for 800 hours, 300 times, across the full temperature range spanned by the three main text datasets ([-1, 30.5] in 0.1-degree increments). To mimic natural communities, rRNA copy numbers ranged from 1 to 10 and were drawn from a geometric distribution, (1 – *p*)^*k*1^ *p*, where *k* is the copy number and the parameter *p* represents the fraction of taxa with copy number = 1 in the starting distribution of the community. We sampled a range of values of the parameter *p* ([0.6, 0.95] in increments of 0.0025). We set most parameters to values midway through the ranges described above (activation energy E = 0.33 eV, prefactor a = 170,000, mean interaction *α_ij_* = 0.5 (with SD = 0.25)). To include temperatures between 0.1-degree increments, we then used a three-degree polynomial fit. In order to fit a value of *p* for each dataset, we first selected a death rate that produced the minimum root-mean-square error across all datasets. We simulated the model at multiple death rates (spanning between 0.03/hr and 0.09/hr) and found that 0.04/hr produced the best fit. To find the best fit, we bootstrapped 1000 trials per dataset, sampled randomly with replacement with sample size equal to that of the dataset. For each sampled data point, we calculated the square of the difference between the observed and the temperature-dependent simulated mean copy numbers (MCN) across all values of μ, and selected the value of *p* for each trial that minimized the root mean square error. We averaged the values of *p* selected for the 1000 trials to obtain the best fit of *p* for each dataset (LMO: *p* = 0.6955 +/− 0.003, mean minimum error = 0.2546; ANT: *p* = 0.7389 +/− 0.0054, mean minimum error = 0.1656; TARA: *p* = 0.8055 +/− 0.0095, mean minimum error = 0.1998).

### Statistical analyses

We used generalized additive models (GAMs(*73*)) to assess the effect of environmental variables on MCN, due to their ability of fitting both linear and non-linear effects and their suitability for modelling large scale trends(*74*). We used the R library “mgcv”(*75*) to construct and fit all GAMs. Smooth terms, “s”, were modeled as a thin plate regression splines(*76*). In the case of TARA ocean GAMs, we used the analogue of thin plate splines for the sphere (two dimensional splines) to model latitude and longitude(*77*). The other parameters were set on default mode. In all instances, we first fitted a full GAM model including the parametric effects of all available environmental variables and the proxy for the environmental gradient (either month, latitude, or depth) as a smooth term. The performance of the full model was then checked by inspecting the distribution of residuals generated via the *gam.check* function in the “performance” package(*78*) and the collinearity among variables estimated using the *check_collinearity* function. Variables with variance inflation factor (VIF) larger than 10 were excluded from the final model. Note that this doesn’t mean that the predictor has no effect on the response variable but rather that its effect is already captured by another predictor included in the model(*79*). For datasets including multiple sampling depths or stations ( ANT, SPOT, P15 GO-SHIP, and TARA), we fitted generalized additive mixed models (GAMMs(*73*)), which allow to specify random effects. The sampling depth or station was, in these cases, included as a random effect to account for related changes in the intercept of the model(*73*).

## Acknowledgments

We thank Dr. Daniel Lundin of Linnaeus University for bioinformatical analysis of the LMO dataset. We also thank Dr. Alma Parada for data from the SPOT time series, Drs. Dana Hunt and Christopher Ward for data from the PICO time series, Dr. Adrià Auladell Martín for data from the SOLA time series, and Dr. Eric Raes for data from the P15 GO-SHIP transect. We thank Dr. JL Weissman for assistance with growth rate estimates using the EGGO database. We thank the members of the Gore and Petrov labs, as well as Dr. Tadashi Fukami, for helpful discussions about the manuscript. This work was funded by Simons Collaboration Principles of Microbial Ecosystems (PriME) award number 54239 and National Institutes of Health grant R01-GM102311 to Jeff Gore.

