## Supplementary Materials for "Warmer temperatures favor slower-growing bacteria in natural marine communities"

**This file includes:**

Supplementary Text
Figs. S1 to S14
Table S1

**Supplementary Text**

Lotka-Volterra pairwise model predicts that increasing temperature favors the slower grower

In this paper, we rely on previous theoretical and experimental results in which we showed that increasing temperature generally favored slower-growing species in pairwise competitions. To explain this result mathematically, we employed the two-species Lotka-Volterra (LV) interspecific competition model:

$$\frac{\dot{N}_i}{N_i} = r_i\left(1 - N_i - \alpha_{ij}N_j\right) \tag{1}$$

where $N_i$ represents the abundance of species $i$ normalized by its carrying capacity, $r_i$ its maximal growth rate, and $\alpha_{ij}$ is a dimensionless competition coefficient quantifying the inhibition of species $i$ by species $j$. The outcomes of the model are completely determined by the competition coefficients: when both $\alpha_{ij} < 1, \alpha_{ji} < 1$, both species coexist; when $\alpha_{ij} < 1$ but $\alpha_{ji} > 1$, species $i$ drives species $j$ to extinction (and vice versa); when both $\alpha_{ij} > 1, \alpha_{ji} > 1$, the result is bistability, in which both species can drive each other extinct, and the winner depends on the initial relative fraction. Thus, the model does not depend on the growth rates—a strongly competing slow grower can drive a fast grower extinct.

Many microbial communities experience mortality that is not driven by competition and which affects the entire community. Importantly, this is true of all laboratory cultures, where cells are removed from the community either continuously (as in a chemostat or turbidostat) or at discrete intervals (as in batch culture). It may also result from predation by bacterivores, or, in the case of our gut microbiota, from waste passing through. Equation (1) can therefore be made more realistic by the introduction of a community-wide mortality rate ($\delta$):

$$\frac{\dot{N}_i}{N_i} = r_i\left(1 - N_i - \alpha_{ij}N_j\right) - \delta \tag{2}$$

Equation (2) can then be re-parameterized back into is original form, such that the outcome is determined completely by the re-parameterized competition coefficients $\hat{\alpha}_{ij}$:

$$\frac{\dot{\hat{N}}_i}{\hat{N}_i} = \hat{r}_i\left(1 - \hat{N}_i - \hat{\alpha}_{ij}\hat{N}_j\right) \tag{3}$$

$$\hat{\alpha}_{ij} = \alpha_{ij}\frac{\left(1 - \frac{\delta}{r_j}\right)}{\left(1 - \frac{\delta}{r_i}\right)} \tag{4}$$

Since $\hat{\alpha}_{ij}$ is a function of growth and death rates, the outcome will shift along with these rates. For example, the faster grower is favored by an increasing death rate. An increasing temperature, on the other hand, will affect growth rather than death. If we assume that maximal growth rates $r_i(T)$ are the only parameters affected by changes in temperature, we can take the derivative of $\hat{\alpha}_{ij}$ with respect to temperature to see which species will benefit from an increase in temperature:

$$\frac{\partial}{\partial T}\hat{\alpha}_{fs} = \alpha_{fs}\frac{\partial}{\partial T}\left(\frac{1-\frac{\delta}{r_s(T)}}{1-\frac{\delta}{r_f(T)}}\right) \tag{5}$$

Here we have used indices $f$ and $s$ to denote the fast and slow grower, respectively. We must choose a model for $r(T)$ in order to continue. As discussed in the main text, we use the Arrhenius equation:

$$r(T) = ae^{-\frac{E_a}{k_B T}} \tag{6}$$

where $a$ is a pre-factor with dimensions of 1/time, $E_a$ is the activation energy (in units of eV), $k_B$ is the Boltzmann constant, and $T$ is the temperature (in Kelvin). In the following equations, we will specify the fast grower with $r_f = a_f e^{-\frac{E_f}{k_B T}}$ and similarly for the slow grower. Plugging this form into Equation 5, we find:

$$\frac{\partial}{\partial T}\hat{\alpha}_{fs} = \alpha_{fs}\frac{\frac{E_s}{k_B}\delta e^{\frac{E_s}{k_B T}}}{a_s T^2\left(1-\frac{\delta}{a_f e^{-\frac{E_f}{k_B T}}}\right)} - \frac{\frac{E_f}{k_B}\delta e^{\frac{E_f}{k_B T}}\left(1-\frac{\delta}{a_s e^{-\frac{E_s}{k_B T}}}\right)}{a_f T^2\left(1-\frac{\delta}{a_f e^{-\frac{E_f}{k_B T}}}\right)^2} \tag{7}$$

For the slow-grower to be favored, the above term should be greater than zero, because this would mean that inhibition of the fast grower by the slow grower increases with temperature (and inhibition of the slow grower by the fast grower decreases):

$$\frac{\frac{E_s}{k_B}\delta e^{\frac{E_s}{k_B T}}}{a_s T^2\left(1-\frac{\delta}{a_f e^{-\frac{E_f}{k_B T}}}\right)} > \frac{\frac{E_f}{k_B}\delta e^{\frac{E_f}{k_B T}}\left(1-\frac{\delta}{a_s e^{-\frac{E_s}{k_B T}}}\right)}{a_f T^2\left(1-\frac{\delta}{a_f e^{-\frac{E_f}{k_B T}}}\right)^2} \tag{8}$$

We can simplify this expression by plugging in $r_f$ and $r_s$ where they appear:

$$\frac{E_s}{r_s T^2 \left(1 - \frac{\delta}{r_f}\right)} > \frac{E_f \left(1 - \frac{\delta}{r_s}\right)}{r_f T^2 \left(1 - \frac{\delta}{r_f}\right)^2} \tag{9}$$

Ultimately, the expression simplifies to a simple relation showing that the slow grower is always favored as temperature increases when the activation energies of the two species are the same (i.e. when $E_s = E_f$):

$$\frac{E_s}{E_f} > \frac{r_s - \delta}{r_f - \delta} \tag{10}$$

In the main text, we assumed equal activation energies of all species, and that differences in growth rates arise from differing rRNA copy numbers (which were absorbed into the pre-factor $a$). Thus, the inequality is always true under our assumptions.

We can repeat this calculation with another model for the dependence of growth rate on temperature. The Ratkowsky model is consistently the best fit for microbial data:

$$r(T) = b^2 (T - T_o)^2 \tag{11}$$

Where $b$ and $T_o$ must be determined by fitting data for a particular species. Repeating the calculation in the same way as above, we find a similar inequality that determines when the slow grower is favored by increasing temperature:

$$\frac{(T - T_{of})}{(T - T_{os})} > \frac{r_s - \delta}{r_f - \delta} \tag{12}$$

When the growth curves do not cross, we can assume that $T_{os} > T_{of}$ and $r_f > r_s$. These assumptions make the left side of the inequality greater than one, while the right side is less than one. Thus, the inequality is always true for a competition between a consistent slow grower and a consistent fast grower, and the LV model predicts that an increasing temperature will always favor a slower grower, provided that the slower grower does not become relatively faster at high temperature.

**Fig. S1.**

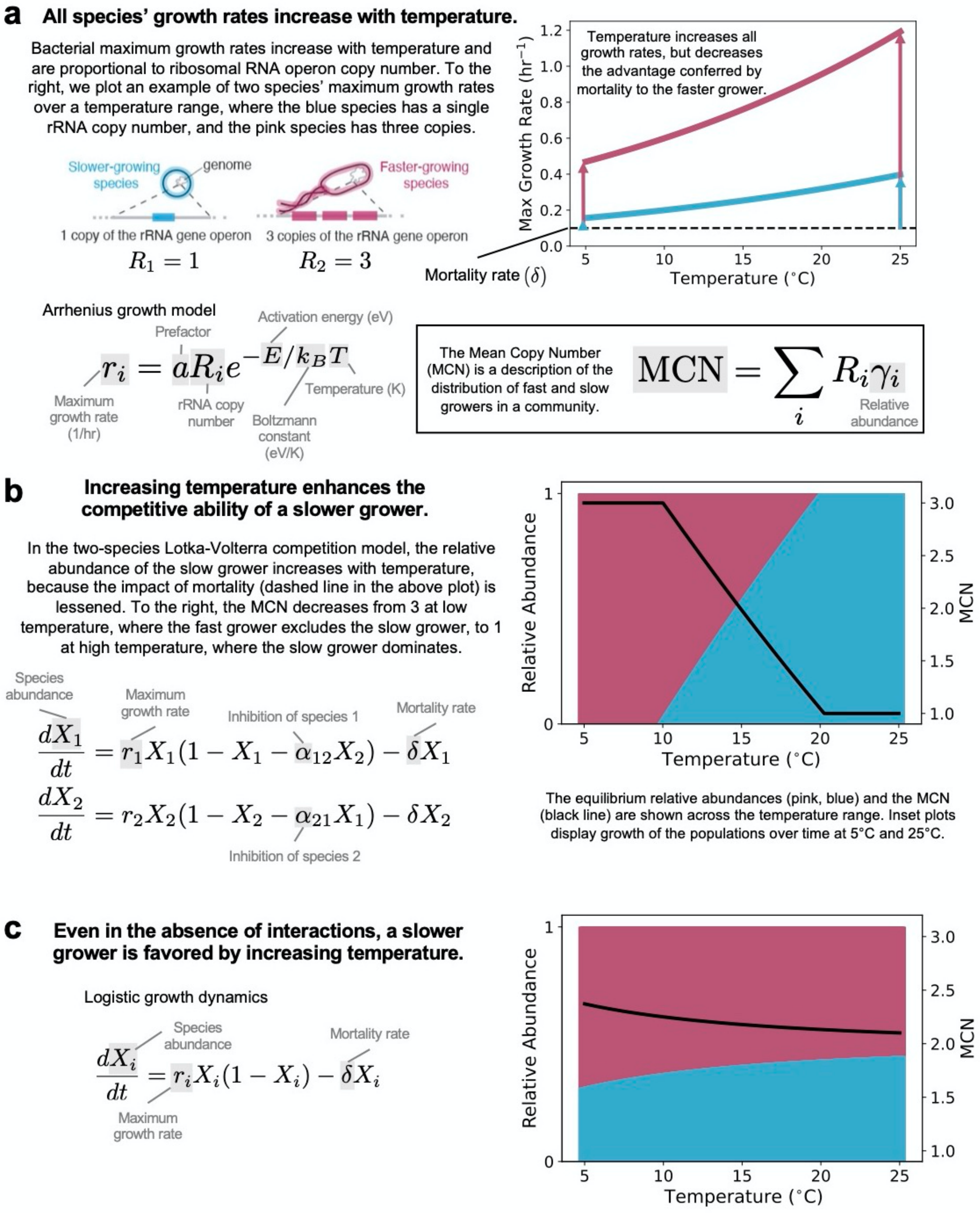


**Two-species Lotka-Volterra competition model predicts that increasing temperature favors the slower grower.** The analytical solutions are shown for the case in which $r = aRe^{-\frac{G}{T}}$, where $G = 3864$ and $a = 1.7e5$ for both species, and $R = 1$ for the blue species and $R = 3$ for the pink species.

**Fig. S2.**

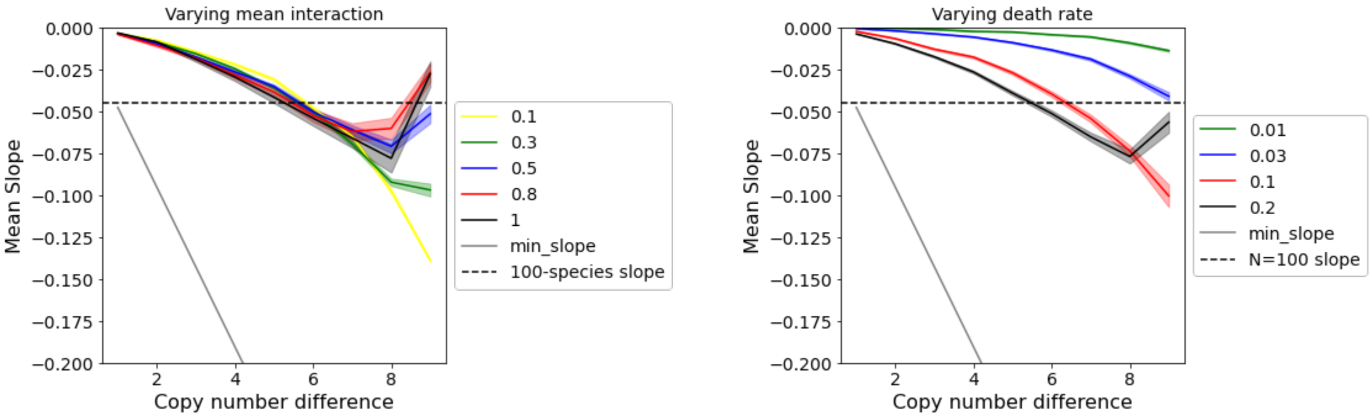


**Two-species Lotka-Volterra solution is independent of mean interaction coefficient, as seen in the left plot.** (Noise on the right end of the plot reflects extreme differences in copy number between the two species leading to fewer instances of coexistence and shallower MCN-temperature slopes.) The two-species solution is sensitive to death rate, with increasing death rate leading to steeper MCN-temperature slopes.

**Fig. S3.**

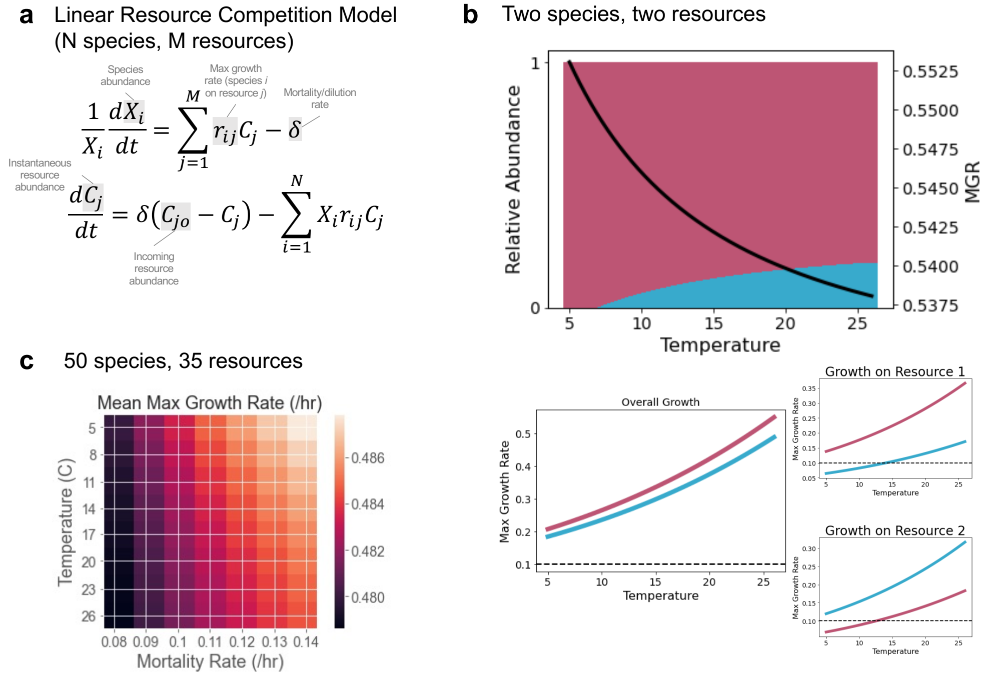


$$\frac{1}{X_i}\frac{dX_i}{dt} = \sum_{j=1}^{M} r_{ij}C_j - \delta$$

$$\frac{dC_j}{dt} = \delta\left(C_{jo} - C_j\right) - \sum_{i=1}^{N} X_i r_{ij} C_j$$

**Consumer resource model predicts that warmer temperatures favor slower growers.** In a resource-explicit model (panel **a**), a species' overall maximum growth rate is additively composed of its maximum growth rates on individual resources. In a two-species, two-resource competition (panel **b**), the overall slower grower benefits from increasing temperature (upper plot), because the relative abundance of the slow grower (blue) increases with temperature and the relative abundance of the fast grower (pink) decreases, causing the abundance-weighted mean growth rate (MGR) to decrease. The growth rates of each species on each resource are equal to $ae^{-\frac{E}{k_B T}}$, where activation energy $E = 0.33\ eV$ for both species on both resources, $a_{11} = 7 * 10^4$, $a_{12} = 1.3 * 10^5$, $a_{21} = 1.5 * 10^5$, and $a_{22} = 7.5 * 10^4$. Mortality/dilution rate $\delta$ was set to 0.1/hr, and indicated by the dashed line in panel **b** (lower plots). The analytical solution is shown. In a competition with 50 species and 35 resources (panel **c**), the MGR describes the distribution of fast and slow growers in the community. The average MGR of 100 simulations is shown for a range of temperatures and mortality/dilution rates. Growth rates on each resource are equal to $ae^{-\frac{G}{T}}$, where $G = 3864$ for all species on all resources, and $a$ was randomly drawn from a uniform distribution, $[5 * 10^4, 3 * 10^5]$. To compute the MGR, the maximum growth rate for each species was summed across all resources (the incoming resource concentration $C_o$ was set to 1 for all resources, so no weighting over resources was necessary), and the relative abundances of species after 500 hours of simulation were used to weight the mean maximum growth rate. Mortality/dilution rate was set to 0.1. As in the Lotka-Volterra model, increasing mortality and decreasing temperature favor fast growers with higher maximum growth rates, while slow growers benefit from decreasing mortality and increasing temperature.

**Fig. S4.**

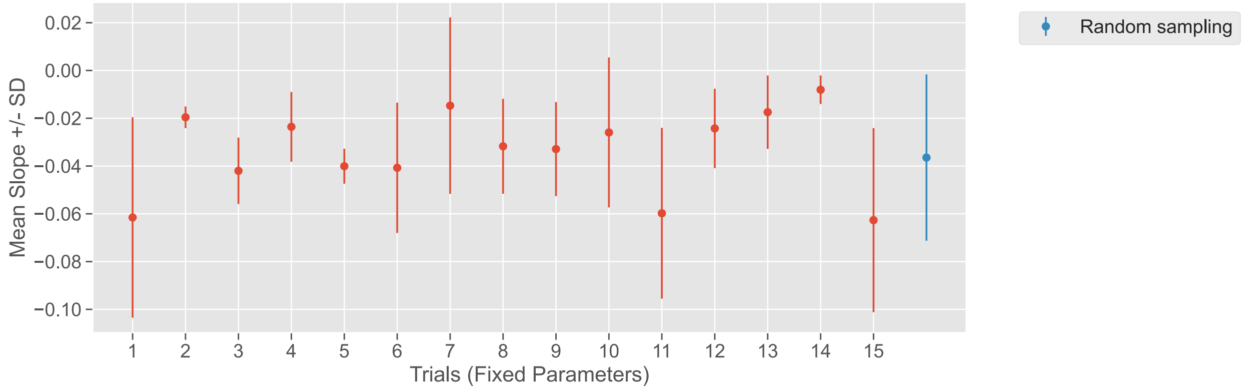


**The mean and standard deviation of MCN-temperature slopes resulting from constant parameter simulations are similar to the mean and standard deviation of simulations performed by randomly sampling parameters.** Red points represent randomly chosen parameters that remained fixed over 500 simulations (interaction coefficient matrices $\alpha_{ij}$ are unique for each simulation, even if mean interaction is fixed). The blue point on the right represents 500 simulations in which parameters were randomly sampled at each simulation (the mean of the normal distribution of $\alpha_{ij}$ was randomly drawn from a uniform distribution [0.1, 1] and the standard deviation was set to half the mean; the geometric distribution parameter $p$ of the rRNA copy numbers was drawn from a uniform distribution [0.6, 0.9]; the mortality rate δ was drawn from a uniform distribution [0.03, 0.2]; the activation energy $E$ was drawn from a uniform distribution [0.1 eV, 0.6 eV]).

**Fig. S5.**

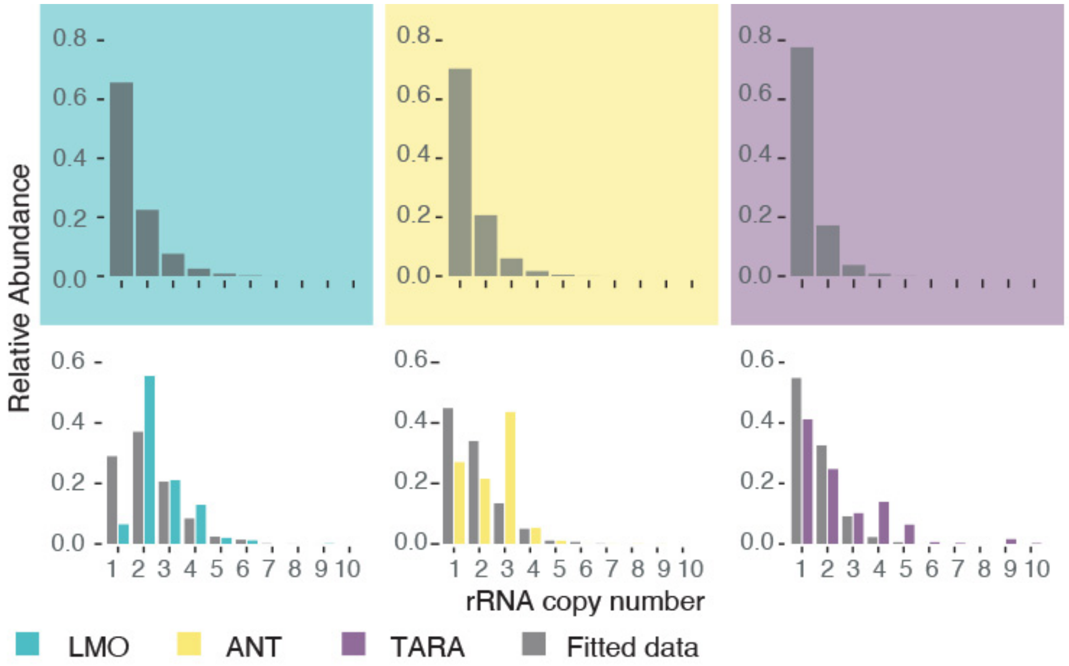


**The final abundance distribution of copy numbers obtained from fitting the GLV model to data resembles the observed one in the three main datasets.** Upper panels: abundance distribution of copy numbers the initialized communities in the model. Lower panels: abundance distribution of the equilibrated communities (grey) compared to the observed distribution of copy numbers in the three main datasets (green, LMO; yellow, ANT; purple, TARA).

**Fig. S6.**

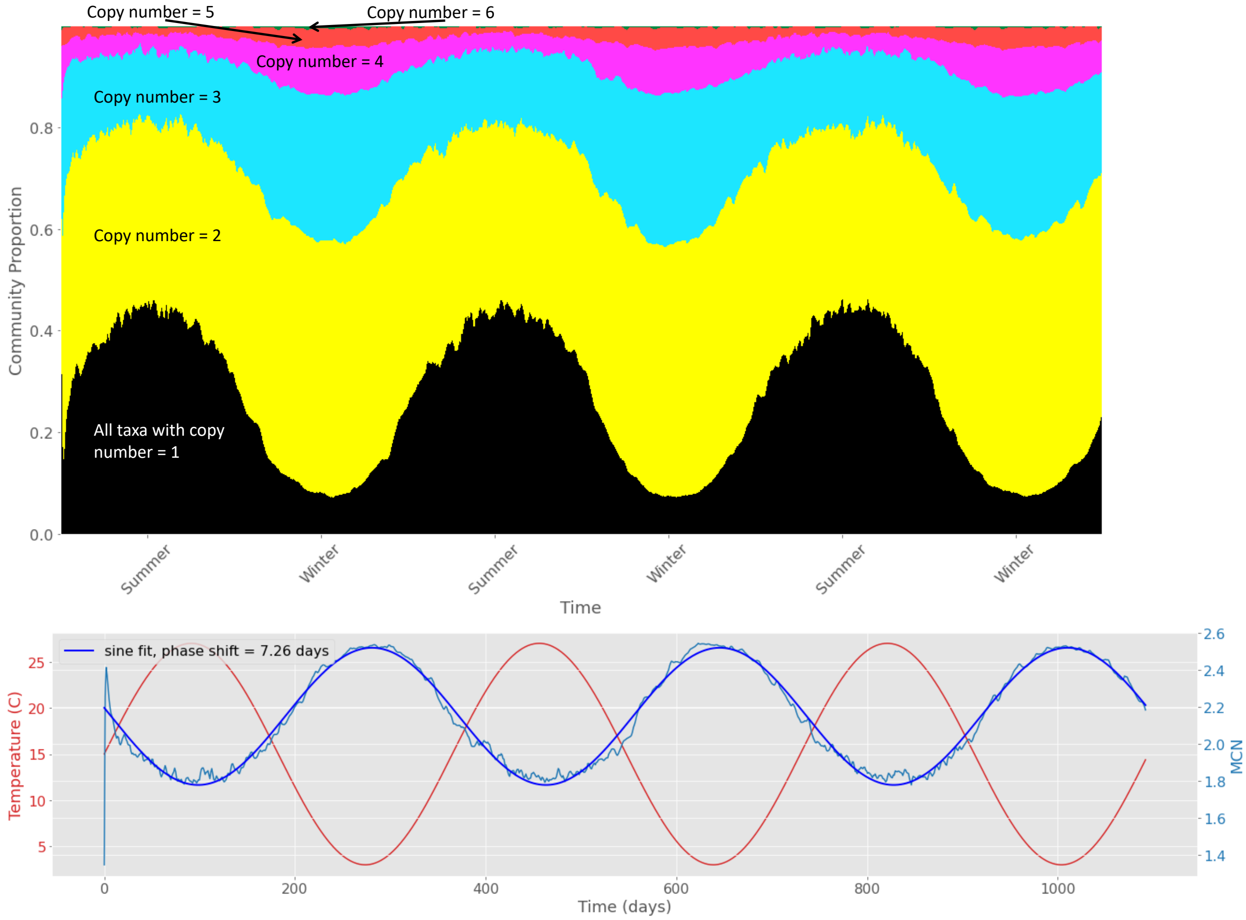


**A continuous-time simulation with sinusoidally oscillating temperature shows that mean copy number (MCN) oscillations lag behind temperature oscillations, resulting in a frequency equal to that of temperature but with a phase shift of about one week (7.26 days) when the MCN was fit to a sinusoidal curve.** The average of 50 Lotka-Volterra simulations is plotted, each with 100 species and mean interaction $\alpha = 0.5$ ($\sigma = 0.25$), mortality rate $\delta = 0.07/hr$, with copy numbers drawn from a geometric distribution, $p(1-p)^{R-1}$, where copy numbers $R$ range from 1 to 10 and $p = 0.75$. Maximum growth rates are equal to $aRe^{-\frac{G}{T}}$, where $R$ is the copy number, $a$ is a prefactor equal to 170,000, the activation energy $G = 3864$, and temperature ranges from 3 to $27°C$. The simulations began with all species at equal relative abundance, and every four days, an abundance of $10^{-4}$ of all species (in units of fraction of single-species carrying capacities in the model) was added to the pool to represent migration and to prevent extinction.

**Fig. S7.**

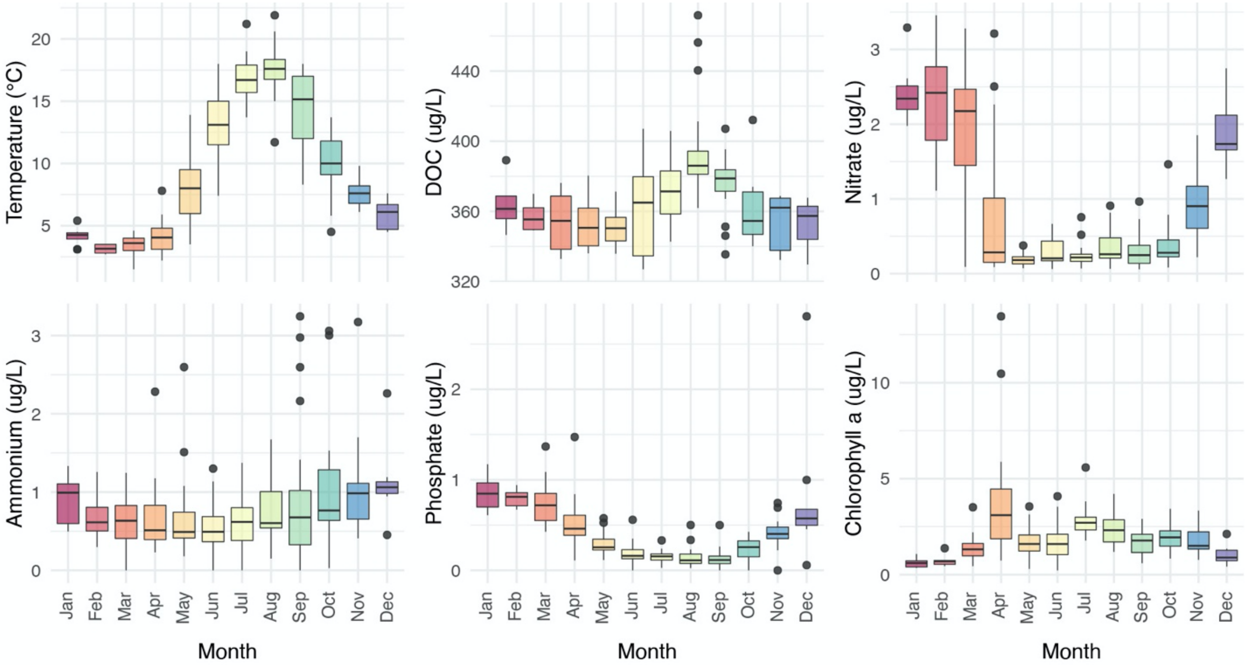


**Many environmental variables show seasonality at the LMO station.** Boxplots showing monthly variation in the environmental variables collected at the LMO station in 8 years (2011-2018).

**Fig. S8.**

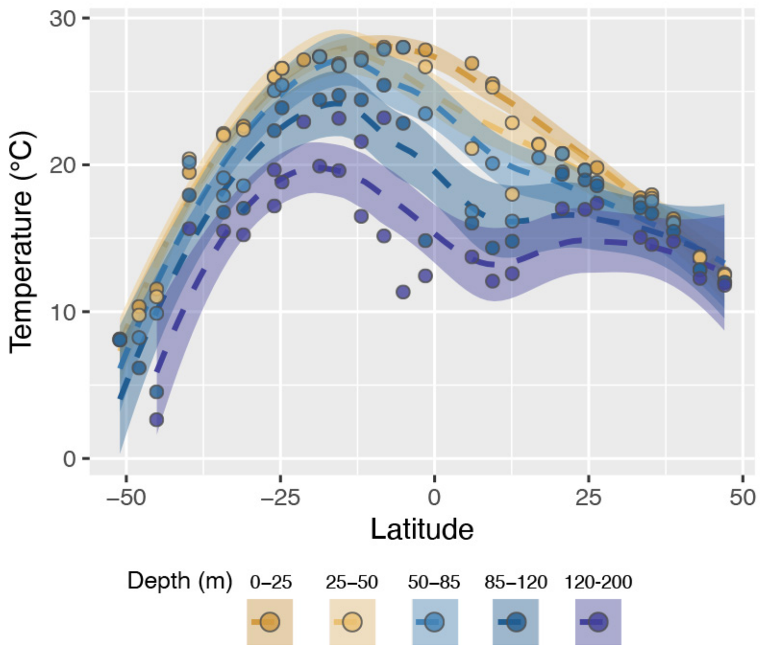


**Temperature profiles at different depths show similar latitudinal trends at ANT sampling stations.** Temperature variations across latitudes in the ANT dataset (dots, observed values, dashed lines: smoothing splines). Colors represent the depth range at which the sample was taken (five within the surface and the 200m).

**Fig. S9.**

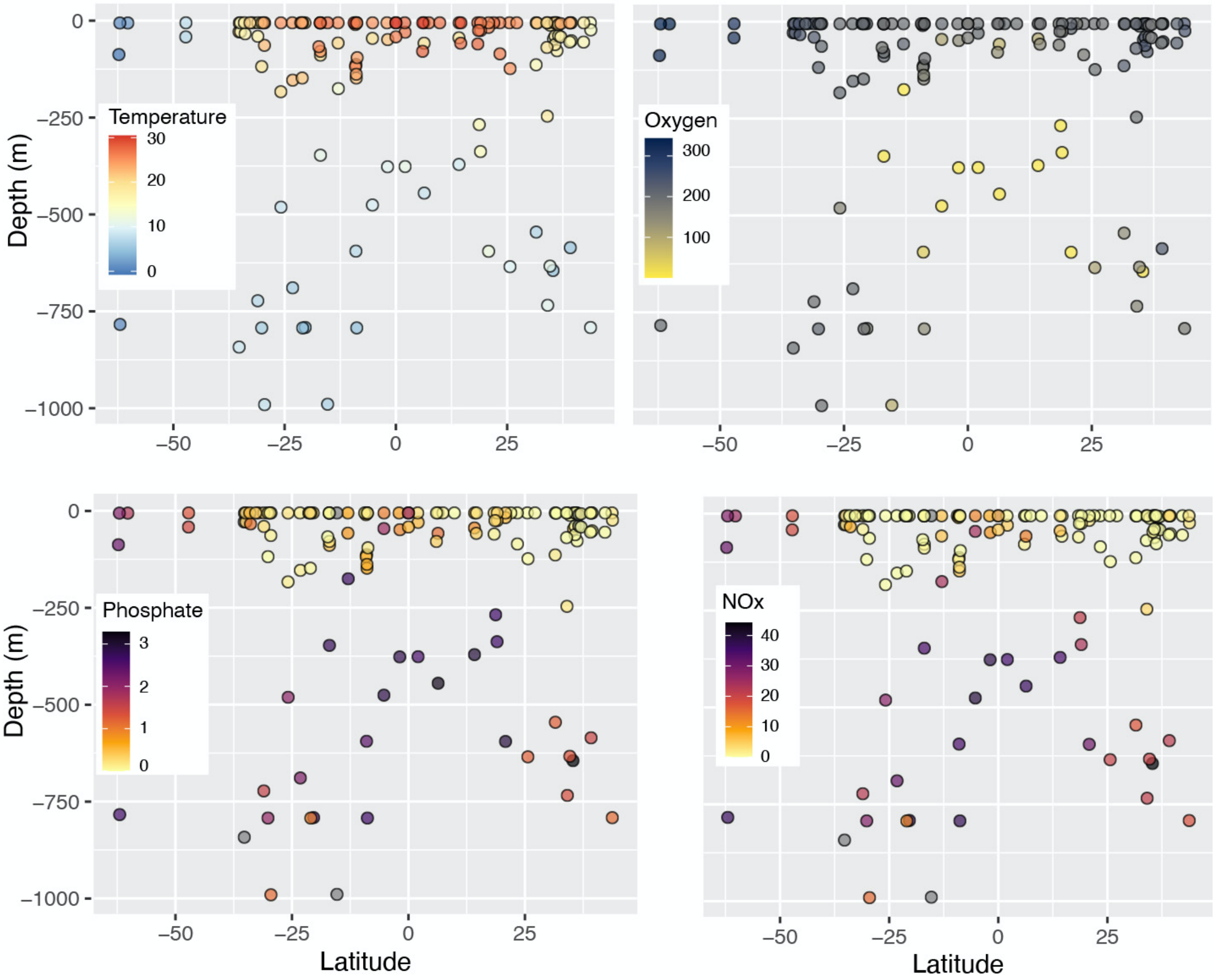


**Temperature and oxygen decrease at high latitudes and with depth, while inorganic nutrients show the opposite pattern.** Observed Temperatures, Oxygen, Phosphate and Nitrate + Nitrite (NOx) concentrations across latitudes and depths sampled in the TARA Ocean project.

**Fig. S10.**

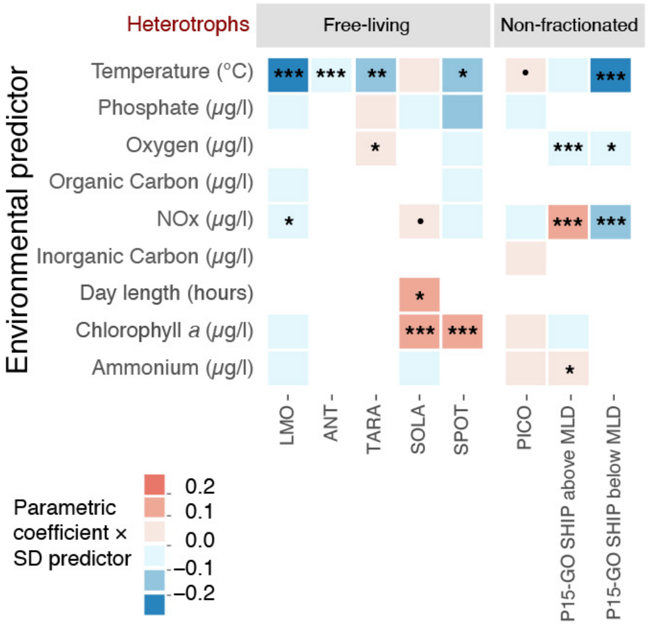


**Considering the effect of primary productivity does not affect the negative effect of temperature on the distribution of fast and slow growing heterotrophs.** Chlorophyll *a* concentration is a proxy for primary productivity and a known driver of several bacterial species(*6*, *76*). Since phototrophic bacteria contain chlorophyll *a*, they must be excluded from the calculation of the MCN to show that the temperature effect is robust to the inclusion of chlorophyll *a* concentration in the multiparametric analyses, which we do for the LMO and SPOT datasets (table S1, but see the results for SOLA time series in table S1). The figure shows a summary of the results of the multi-parametric regression on MCN of heterotrophic communities of all the datasets included in the study (free-living and non-fractionated samples). Tile colors represent the magnitude and sign of the parametric coefficient estimated for each available environmental variable multiplied by its standard deviation. Statistical significance of the parametric coefficients estimated through the models is indicated by symbols: '***' 0.001 '**' 0.01 '*' 0.05 '.' 0.1 ' ' 1.

**Fig. S11.**

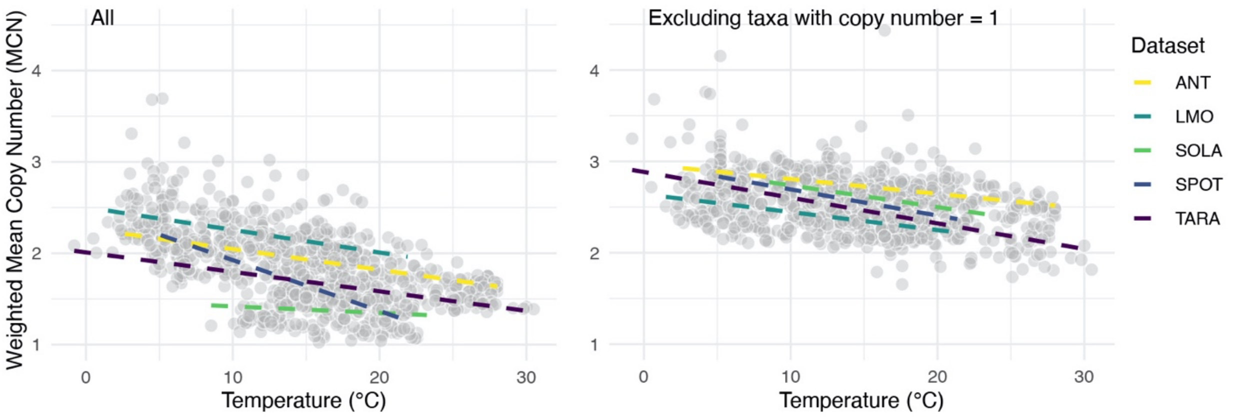


**The relative abundance of free-living slow-growers increases with temperature, regardless of whether oligotrophs are included in the estimation of the weighted mean copy number (MCN) or excluded from it.** Left panel: MCN calculated on all taxa; right panel: MCN calculated excluding taxa with copy number equal to one. Colored dashed lines represent linear regressions between temperature and MCN for each dataset. The relationship between MCN and temperature is negative and statistically significantly for LMO (temperature sensitivity = -0.003 $\pm$ 0.001 ΔMCN/°C, $p = 0.0005$***), ANT (temperature sensitivity = -0.023 $\pm$ 0.002 ΔMCN/°C, $p = 6.83e\text{-}15$), and TARA (temperature sensitivity = -0.021 $\pm$ 0.002 ΔMCN/°C, $p = 1.08e\text{-}12$***), SPOT (temperature sensitivity = -0.056 $\pm$ 0.003 ΔMCN/°C, $p < 2e\text{-}16$***), while it is not significant in SOLA (temperature sensitivity = -0.007 $\pm$ 0.006 ΔMCN/°C, $p = 0.273$ n.s.). The relationship between MCN calculated excluding taxa with copy number = 1 and temperature is negative and statistically significantly for all datasets: LMO temperature sensitivity = -0.02 $\pm$ 0.003 ΔMCN/°C, $p = 6.77e\text{-}12$***, ANT temperature sensitivity = -0.016 $\pm$ 0.002 ΔMCN/°C, $p = 1.1e\text{-}13$***, TARA temperature sensitivity = -0.028 $\pm$ 0.003 ΔMCN/°C, $p < 2e\text{-}16$***, SOLA temperature sensitivity = -0.024 $\pm$ 0.001 ΔMCN/°C, $p = 1.82e\text{-}09$***, SPOT temperature sensitivity = -0.03 $\pm$ 0.003 ΔMCN/°C, $p < 2e\text{-}16$***.

**Fig. S12.**

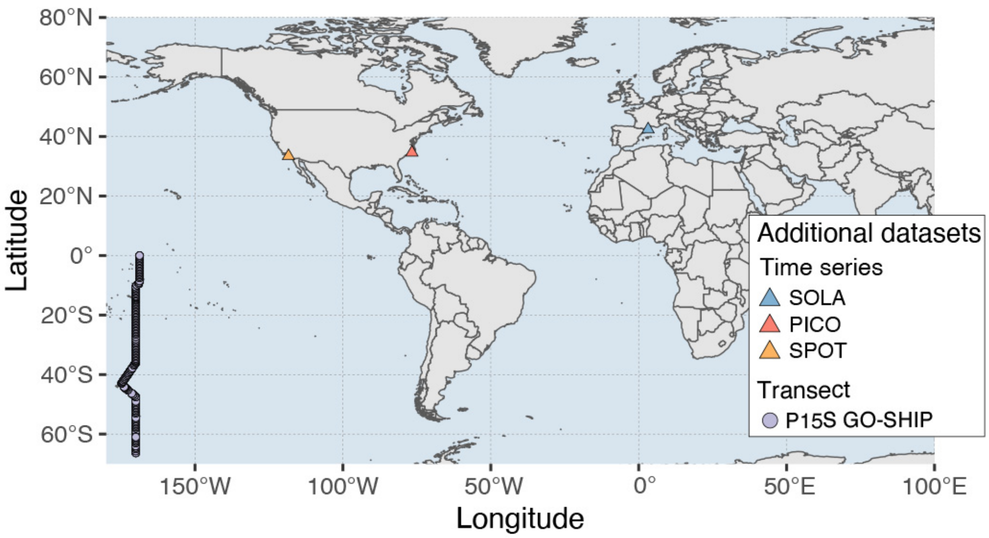


**We considered a total of 7 datasets reporting the composition of marine bacterial communities.** Map of the additional datasets included in the study.

**Fig. S13.**

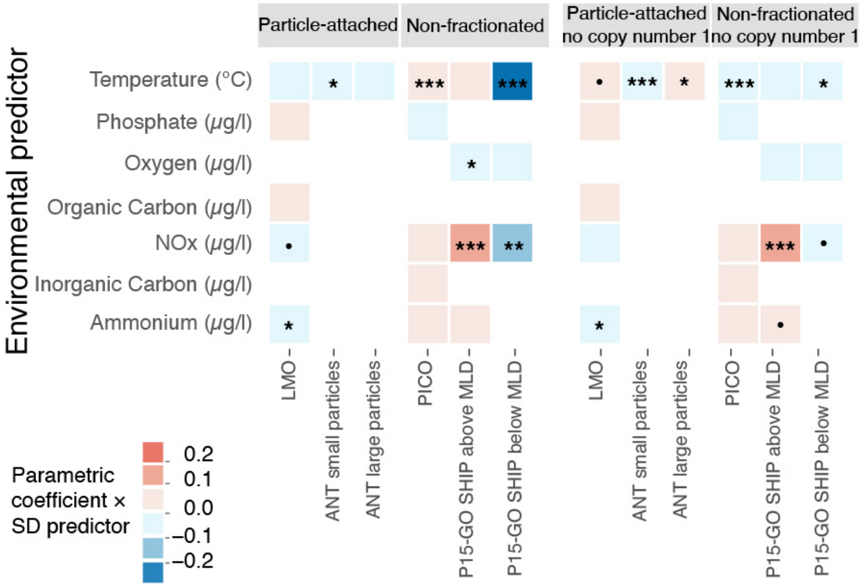


**The effects of temperature on the distribution of fast and slow growers attached to particles are idiosyncratic.** Summary of the results of the multi-parametric regression on MCN of particle-attached (available for LMO timeseries and ANT transect) and non-fractionated (available for PICO time series and P15 GO-SHIP transect) communities. Tile colors represent the magnitude and sign of the parametric coefficient estimated for each available environmental variable multiplied by its standard deviation. Statistical significance of the parametric coefficients estimated through the models is indicated by symbols: '***' 0.001 '**' 0.01 '*' 0.05 '.' 0.1 ' ' 1.

**Fig. S14.**

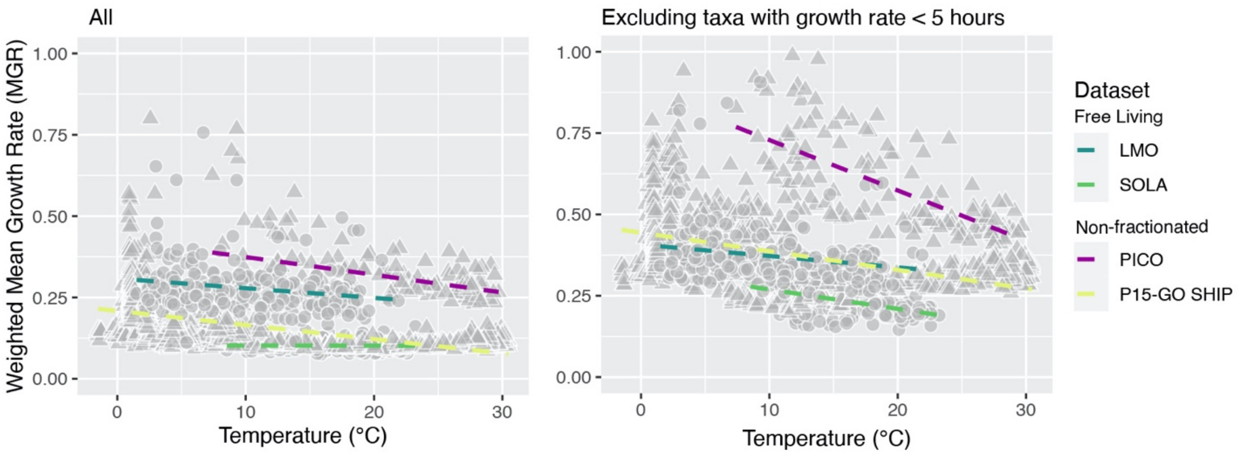


**Figure S14. Estimating the distribution of fast and slow growing taxa with an alternative method yields a negative relationship with temperature.** To test whether our conclusions depend on the method for estimating growth rates with rRNA copy number, we used another method based on codon usage bias (48) (this method is optimized for metagenomic datasets, although a 16S reference is available). The relationship between MGR (weighted mean growth rate, obtained from codon usage bias estimations) and temperature is negative and statistically significantly for LMO (temperature sensitivity = - 0.003±0.001 ΔMCN/°C, $p = 0.0005$***), PICO (temperature sensitivity = -0.005±0.001 ΔMCN/°C, p = 2.56e-07***), and P15 GO-SHIP (temperature sensitivity = -0.004 ± 0.0002 ΔMCN/°C, $p < 2e\text{-}16$ ***), while it is not significant in SOLA (temperature sensitivity = -0.00003 ± 0.0004 ΔMCN/°C, $p = 0.947$ n.s.). The relationship between MGR calculated excluding taxa with growth rate lower than 5 hours (48) and temperature is negative and statistically significantly for all datasets: LMO temperature sensitivity = - 0.004 ± 0.001 ΔMCN/°C, p = 0.0005***, PICO temperature sensitivity = -0.017±0.001 ΔMCN/°C, $p < 2e\text{-}16$***, P15 GO-SHIP temperature sensitivity = -0.006 ± 0.0003 ΔMCN/°C, p < 2e-16***, SOLA temperature sensitivity = -0.006 ± 0.001 ΔMCN/°C, $p = 1.82e\text{-}09$***.

**Table S1.**

Effects of environmental variables on MCN of entire, copiotroph and heterotroph communities of all datasets included in the study. LMO, ANT, TARA, SOLA and SPOT comprise free-living communities, while PICO and P15 GO-SHIP include only non-fractionated samples. For LMO and ANT particle-attached communities are also available. *Organic carbon* is the dissolved fraction (DOC) in LMO and the particulate fraction (POC) in SPOT. *Light* measurements also differ between datasets: in SOLA it is measured as Day length (in hours); in PICO as insolation (kWh $m^{-2}d^{-1}$). Effects of environmental variables on MCN have been assessed using generalized additive (*gam*) or generalized additive mixed models (*gamm*). The latter were used when multiple sampling depths or stations were available the dataset, which were included in the random part of the model (ANT, SPOT, P15 GO-SHIP, and TARA). The intercept of the model and parametric coefficients ($\gamma$) for environmental variables are reported as Estimate (St. Error) Pr(>|t|). Significance codes: '***' 0.001 '**' 0.01 '*' 0.05 '.' 0.1 ' ' 1. Statistically significant parametric coefficients are highlighted in different shades of yellow. Adjusted $R^2$ are included. Smooth terms and random terms are omitted.

| | | *Intercept* | *Temperature* | *Phosphate* | *Ammonium* | *Nitrate* | *Organic Carbon* | *Inorganic Carbon* | *Oxygen* | *Light* | *Chlorophyll a* | *Adj. $R^2$* |
|---|---|---|---|---|---|---|---|---|---|---|---|---|
| All | LMO | 2.966 (0.370) <2e-16 *** | - 0.031 (0.006) 1.82e-07*** | - 0.023 (0.138) 0.868 | - 0.082 (0.033) 0.015 * | - 0.068 (0.039) 0.806 · | - 0.001 (0.001) 0.111 | | | | | 0.259 |
| | ANT | 2.060 (0.078) <2e-16 *** | - 0.011 (0.005) 0.007 ** | | | | | | | | | 0.57 |
| | TARA | 1.766 (0.167) < 2e-16 *** | - 0.018 (0.005) 0.0008 *** | 0.094 (0.047) 0.047 * | | High collinearity (excluded) | | | 0.001 (0.0004) 0.155 | | | 0.691 |
| | SOLA | 0.702 (0.170) <2e-16 *** | - 0.013 (0.007) 0.087 · | -0.388 (0.647) 0.550 | -0.014 (0.080) 0.858 | 0.061 (0.023) 0.011 * | | | | 0.066 (0.012) 0.0609 *** | | 0.177 |
| | SPOT | 2.605 (0.139) < 2e-16 *** | -0.052 (0.011) 6.03e-06 *** | | | | | | - 0.067 (0.213) 0.002 ** | | | 0.401 |
| Copiotro | LMO | 2.621 (0.341) 1.33e-12 *** | - 0.028 (0.006) 8.58e-06 *** | - 0.015 (0.130) 0.910 | - 0.076 (0.031) 0.015 * | - 0.056 (0.036) 0.122 | 0.001 (0.001) 0.555 | | | | | 0.211 |
| | ANT | 2.754 (0.050) | - 0.005 (0.003) 0.082 · | | | | | | | | | 0.563 |

| | | | | | | | | | | | | |
|---|---|---|---|---|---|---|---|---|---|---|---|---|
| phs | | <2e-16 *** | | | | | | | | | | |
| | TARA | 2.104 (0.079) <2e-16 *** | -0.023 (0.009) 0.008 ** | 0.099 (0.080) 0.216 | | High collinearity (excluded) | | | 0.002 (0.001) 0.020 * | | | 0.703 |
| | SOLA | 3.118 (0.174) <2e-16 *** | -0.015 (0.008) 0.056 · | 0.088 (0.662) 0.894 | -0.011 (0.082) 0.892 | 0.007 (0.024) 0.774 | | | | -0.023 (0.013) 0.072 · | | 0.092 |
| | SPOT | 3.028 (0.074) <2e-16 *** | -0.034 (0.009) 0.060 *** | | | | | | 0.006 (0.017) 0.737 | | | 0.308 |
| Heterotrophs | LMO | 2.380 (0.441) 2.41e-07 *** | -0.030 (0.009) 0.0007 *** | 0.271 (0.193) 0.162 | -0.004 (0.043) 0.926 | -0.116 (0.052) 0.027 * | 0.0002 (0.001) 0.851 | | | | 0.017 (0.030) 0.558 | 0.193 |
| | ANT | 2.104 (0.079) <2e-16 *** | -0.015 (0.004) 0.0005 *** | | | | | | | | | 0.601 |
| | TARA | 1.714 (0.165) <2e-16 *** | -0.015 (0.005) 0.005 ** | 0.106 (0.046) 0.025 * | | | | | 0.001 (0.0004) 0.098 · | | | 0.671 |
| | SOLA | 0.102 (0.419) 0.807 | 0.002 (0.014) 0.881 | -0.164 (0.730) 0.823 | -0.007 (0.086) 0.938 | 0.045 (0.026) 0.082 · | | | | 0.084 (0.034) 0.014 * | 0.249 (0.060) 5.39e-05 *** | 0.268 |
| | SPOT | 2.232 (0.350) 8e-07 *** | 0.032 (0.014) 0.034 * | -0.666 (0.375) 0.087 · | | -0.005 (0.036) 0.883 | -0.0002 (0.0003) 0.603 | | -0.018 (0.039) 0.643 | | 0.112 (0.030) 0.0009 *** | 0.469 |
| Particle | LMO | 2.353 (0.52) 1.15e-05 *** | -0.005 (0.009) 0.588 | 0.002 (0.198) 0.993 | -0.111 (0.047) 0.02 * | -0.099 (0.055) 0.073 · | 0.001 (0.001) 0.595 | | | | | 0.041 |
| | ANT small | 2.356 (0.090) <2e-16 *** | -0.010 (0.005) 0.03 * | | | | | | | | | 0.426 |
| | ANT large | 2.213 (0.044) <2e-16 *** | -0.003 (0.002) 0.11 | | | | | | | | | 0 |
| Part | LMO | 2.54 (0.245) <2e-16 *** | -0.014 (0.008) 0.08 · | 0.095 (0.192) 0.62 | -0.113 (0.046) 0.015 * | -0.077 (0.05 | 0.0001 (0.00 | | | | | 0.09 |

| | | | | | | | | | | | |
|---|---|---|---|---|---|---|---|---|---|---|---|
| | | | | | | 3) 0.15 | 1) 0.83 | | | | |
| copio | ANT small | 2.842 (0.06) <2e-16 *** | -0.016 (0.003) 1.29e-06 *** | | | | | | | | 0.242 |
| | ANT large | 2.342 (0.039) <2e-16 *** | 0.004 (0.002) 0.036 * | | | | | | | | 0.015 |
| Non-fract | PICO | 1.32(0.472) 0.006 ** | 0.011 (0.002) 1.78e-05 *** | - 0.344 (0.443) 0.439 | 0 (0) 0.942 | 0.021 (0.119) 0.858 | | 0.0001 (0.0002) 0.618 | | High collinearity (excluded) | 0.127 |
| | P15-GOSHIP above MLD | 1.645 (0.146) <2e-16 *** | 0.002 (0.002) 0.496 | | 0.054 (0.051) 0.286 | 0.010 (0.002) 1.33e-09 *** | | | -0.001 (0.0004) 0.015 * | | 0.364 |
| | P15-GOSHIP below MLD | 2.162 (0.382) 1.17e-15 *** | - 0.082 (0.010) 4.07e-14 *** | | | -0.025 (0.007) 0.0004 *** | | | -0.001 (0.001) 0.032 * | | 0.478 |
| Non-frac copio | PICO | 1.473(0.173) 2.13e-14*** | - 0.007 (0.001) 2.08e-12 *** | -0.085 (0.163) 0.6 | 0 (0) 0.496 | 0.052 (0.043) 0.232 | | 0.0004 (0.0001) *** | | High collinearity (excluded) | 0.315 |
| | P15-GOSHIP above MLD | 2.412 (0.168) < 2e-16 *** | - 0.0003 (0.003) 0.917 | | 0.110 (0.058) 0.060 · | 0.012 (0.002) 2.92e-10 *** | | | -0.001 (0.0005) 0.246 | | 0.624 |
| | P15-GOSHIP below MLD | 3.650 (0.0470) 4.54e-14 *** | - 0.032 (0.013) 0.013 * | | | -0.016 (0.009) 0.062 · | | | -0.001 (0.0001) 0.120 | | 0.245 |
